# Berberine Bridge enzyme-like oxidases orchestrate homeostatic control and signaling of oligogalacturonides in defense and wounding

**DOI:** 10.1101/2024.06.28.601198

**Authors:** Ascenzo Salvati, Alessandra Diomaiuti, Federica Locci, Matteo Gravino, Giovanna Gramegna, Muhammad Ilyas, Manuel Benedetti, Sara Costantini, Monica De Caroli, Baptiste Castel, Jonathan D. G. Jones, Felice Cervone, Daniela Pontiggia, Giulia De Lorenzo

**Affiliations:** Department of Biology and biotechnologies ’Charles Darwin’ Sapienza University of Rome, 00185 Rome, Italy; Department of Biological and Environmental Sciences and Technologies, University of Salento, Campus Ecotekne, 73100 Lecce, Italy; The Sainsbury Laboratory, University of East Anglia, Norwich Research Park, Colney Lane, Norwich, NR4 7UH, UK

## Abstract

Plant immunity is triggered by endogenous elicitors known as damage-associated molecular patterns (DAMPs). Oligogalacturonides (OGs) are DAMPs released from the cell wall (CW) demethylated homogalacturonan during microbial colonization, mechanical or pest-provoked tissue damage, and physiological CW remodeling. Berberine Bridge Enzyme-like (BBE-l) proteins named OG oxidases (OGOXs) have been proposed to govern OGs homeostasis, which is necessary to avoid deleterious growth-affecting hyper-immunity and possible cell death. Using OGOX1 over-expressing lines and *ogox1/2* double mutants, we show that these enzymes determine the levels of active OGs vs. inactive oxidized products (oxOGs). The *ogox1/2*-deficient plants have elevated levels of OGs, while plants overexpressing OGOX1 accumulate oxOGs. The balance between OGs and oxOGs affect disease resistance against *Pseudomonas syringae pv tomato*, *Pectobacterium carotovorum,* and *Botrytis cinerea* depending on the microbial capacity to respond to OGs and metabolize oxOGs. Gene expression upon plant infiltration with OGs reveals that OGOXs orchestrate OG signaling in defense as well as upon tissue damage, pointing to these enzymes as apoplastic players in immunity and tissue repair.

**Teaser:** Oxidases control the homeostasis of oligogalacturonides in the cell wall and play a pivotal role in the plant immunity.

## INTRODUCTION

Plants have evolved, likely before animals, a sophisticated innate immune system to protect themselves against pathogens. During a pathogen attack, the recognition of danger signals occurs through specialized transmembrane receptors known as PRRs (Pattern Recognition Receptors), which recognize a plethora of pathogenic signals known as Microbe-Associated Molecular Patterns (MAMPs) and Damage-Associated Molecular Patterns (DAMPs) (*1–3*).

Oligogalacturonides (OGs) are so far the best characterized DAMPs, deriving from the degradation of the plant cell wall (CW). They are oligomers of alpha-1,4-linked galacturonosyl residues released through the partial hydrolysis of homogalacturonan (HGA) that is a major component of pectin. OGs with a degree of polymerization (DP) between 10 and 15 show the highest immunity-triggering activity (*4–7*) while shorter OGs may have a limited elicitor activity (*8*) or even work as repressors of immunity (*9, 10*). Both pathogen- or plant-secreted enzymes such as polygalacturonases (PGs) may catalyze the formation of OGs upon infection, wounding or physiological cell wall remodeling (*5, 11*). An important notion introduced several year ago is that the release of elicitor-active OGs from HG is not an event due to the casual loss of cell wall integrity but needs the concerted action and specific interaction of PGs with their plant-derived protein inhibitors named PGIPs (*12, 13*). A recent work provides a structural demonstration that mechanistically validates this notion. The interaction between *Phaseolus vulgaris* PGIP2 (*Pv*PGIP2) and *Fusarium phyllophilum* PG (*Fp*PG) creates a substrate binding site on the *Pv*PGIP2-FpPG complex, assembling a new binary enzyme with a boosted substrate binding activity and an altered substrate preference that preferentially produces long elicitor-active OGs vs. short immune-repressive OGs (*10*). The PG-PGIP interaction represents a unique plant mechanism to convert a pathogen virulence activity into a defense trigger and supports the view that OG signaling is an important component in plant immunity.

Analysis of the full-genome expression reveals that OGs influence the expression of ∼4000 genes in *Arabidopsis* (*5*). Accumulation of OGs increases plant resistance against pathogens but their hyper- accumulation results in plant growth penalties (*14*), reflecting the phenomenon known as growth– defense trade-off (*15*). The over-accumulation of OGs can even lead to deleterious hyper-immunity leading to cell death (*14*). A mechanism for controlling the homeostasis of OGs as well as other cell wall derived DAMPs such as cellulose and hemicellulose fragments may rely on specific oligosaccharide oxidases encoded by the Berberine-Bridge Enzyme–like (BBE-l) gene family (*16–18*). The BBE-like protein family comprises 27 members (*19*) and at least four members are capable of oxidizing OGs. These are OGOX1/At4g20830/BBE20, according to the nomenclature reported in (*19*), OGOX2/At4g20840/BBE21, OGOX3/At1g11770/BBE2 and OGOX4/At1g01980/BBE1 (*18*).

Oxidation impairs the DAMP activity of OGs and produces H2O2 as a secondary product. In simultaneous co-treatments, oxidized OGs did not interfere with the ability of OGs of up-regulating the defence-related genes *RetOx* (At1g26380/*BBE3*), also known as *FOX1* (*20*) and *CYP81F2* (At5g57220) (*21*).

Expression of *OGOX1* is up-regulated upon elicitation with OGs or bacterial MAMPs such as flg22 and elf18 (*1*), and upon pathogen infection in a strictly coordinated manner with *CELLOX1*, i.e. a BBE-l protein capable of oxidizing cellulose fragments and mixed linked beta-glucans (*16, 17*). Overexpression of *OGOX1* enhances the oxidation/inactivation of OGs but, nevertheless, enhances the resistance to the necrotroph *Botrytis cinerea (Bc)*, due to the inability of the fungus to efficiently utilize the oxidized OGs as a carbon source (*18*). Whether OGOXs interfere also with OG-mediated signaling in immune responses, however, has never been shown. We have used two bacterial pathogens in addition to *Bc* and, through a reverse genetic approach, we demonstrate that OGOXs play a key role in the homeostasis of OGs in the apoplast and tune the immune responses.

## RESULTS

### OGOXs are apoplastic proteins

Since the DAMP activity of OGs takes place in the apoplast, the proposed role of OGOXs in maintaining the OG homeostasis requires the demonstration that these enzymes are indeed localized in the apoplast. We therefore determined the subcellular localization of OGOX1 and its closest paralog OGOX2. Both enzymes possess a putative N-terminal signal peptide for translocation into the endoplasmic reticulum and are predicted to be extracellular proteins, according to SIGNALP prediction. Both proteins, tagged with GFP at the C-terminus, were transiently expressed in adult tobacco leaves by agroinfiltration and their co-localization with the apoplastic RFP-tagged *Phaseolus vulgaris* polygalacturonase-inhibiting protein (PvPGIP2) (*22*), was analyzed by confocal microscopy, also upon plasmolysis. As a control, CELLOX2, already shown to be apoplastic (*16*), was analyzed in parallel. Both OGOX1 and OGOX2 clearly showed an apoplastic localization (Fig. 1).

**Fig. 1.**
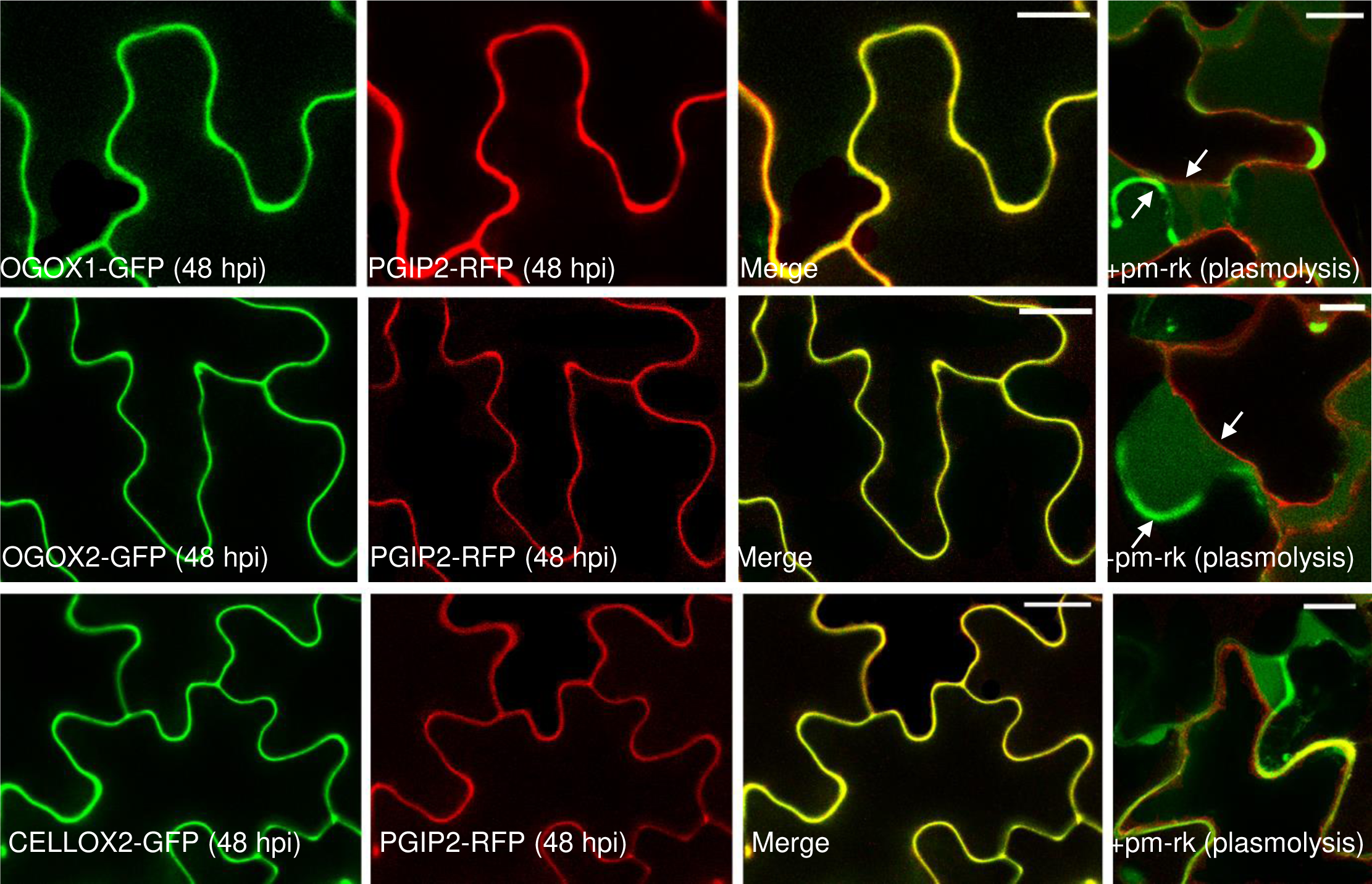
Transient expression of OGOX1-GFP, OGOX2-GFP and CELLOX2-GFP in tobacco epidermal cells. OGOX1-GFP, OGOX2-GFP and CELLOX2-GFP labelled the cell wall, colocalizing with the cell wall marker PGIP2-RFP at 48 h post-infiltration (hpi). Plasmolyzed cells coexpressing OGOX1- GFP, OGOX2-GFP, CELLOX2-GFP and the plasma membrane marker, pm-rk, showed the presence of these proteins in the cell wall, the arrows evidence the green fluorescent cell wall and the retracted red fluorescent plasma membrane. Scale bars 20 µm for OGOX1-GFP. OGOX2-GFP, CELLOX2-GFP and PGIP2-RFP, 10 µm for pm-rk (plasmolysis).

### *OGOX1* is transcriptionally up regulated during infection, wounding and elicitor treatments

The transcriptional regulation of *OGOX1* expression was analyzed in response to infection, wounding and elicitor treatment in adult leaves, using two lines expressing the *GUS* reporter gene under the *OGOX1* promoter (OGOX1::GUS, lines #2.3 and #3.2; see molecular characterization in fig. S1A). *OGOX2* was not analyzed because it is not up-regulated during the immune response (*17*). Upon infection with *Bc*, histochemical staining showed that GUS activity is moderately induced in both OGOX1::GUS lines mainly in the vasculature (Fig. 2A and fig. S1B). Upon infection with bacterial pathogens *Pseudomonas syringae pv tomato* DC3000 (*Pst*) and *Pectobacterium carotovorum* (*Pc),* GUS activity was more vigorously induced in both lines (*Pst*, Fig. 2B and fig. S1C; *Pc*, Fig. 2C and fig. S1D). In the case of infection with *Pst,* there is a diffused and spread induction. Whereas in the case of *Pc*, staining was observed around the damaged tissue and also at the cut site of the petiole in both infected and uninfected excised leaves with some staining spreading to the proximal vasculature (Fig. 2D and fig. S1E). Staining was also observed at the needle-punctured uninfected sites (Fig. 2C and fig. S1D), indicating an up-regulation of *OGOX1* not only upon infection but also in response to mechanical damage. This was confirmed by the observed marked increase of GUS staining around the damaged tissue in leaves at 1 h after crushing (Fig. 2E and fig. S1F). The response of both lines was also examined upon infiltration with either OGs or the MAMP flg22. Leaves showed an intense GUS staining on the entire leaf area, including the vasculature, at 1-hour post-treatment, compared to control leaves infiltrated with water (Fig. 2F and fig. S1G). Overall, these data indicate that *OGOX1* expression is induced upon infection, wounding and treatment with elicitors.

**Fig. 2.**
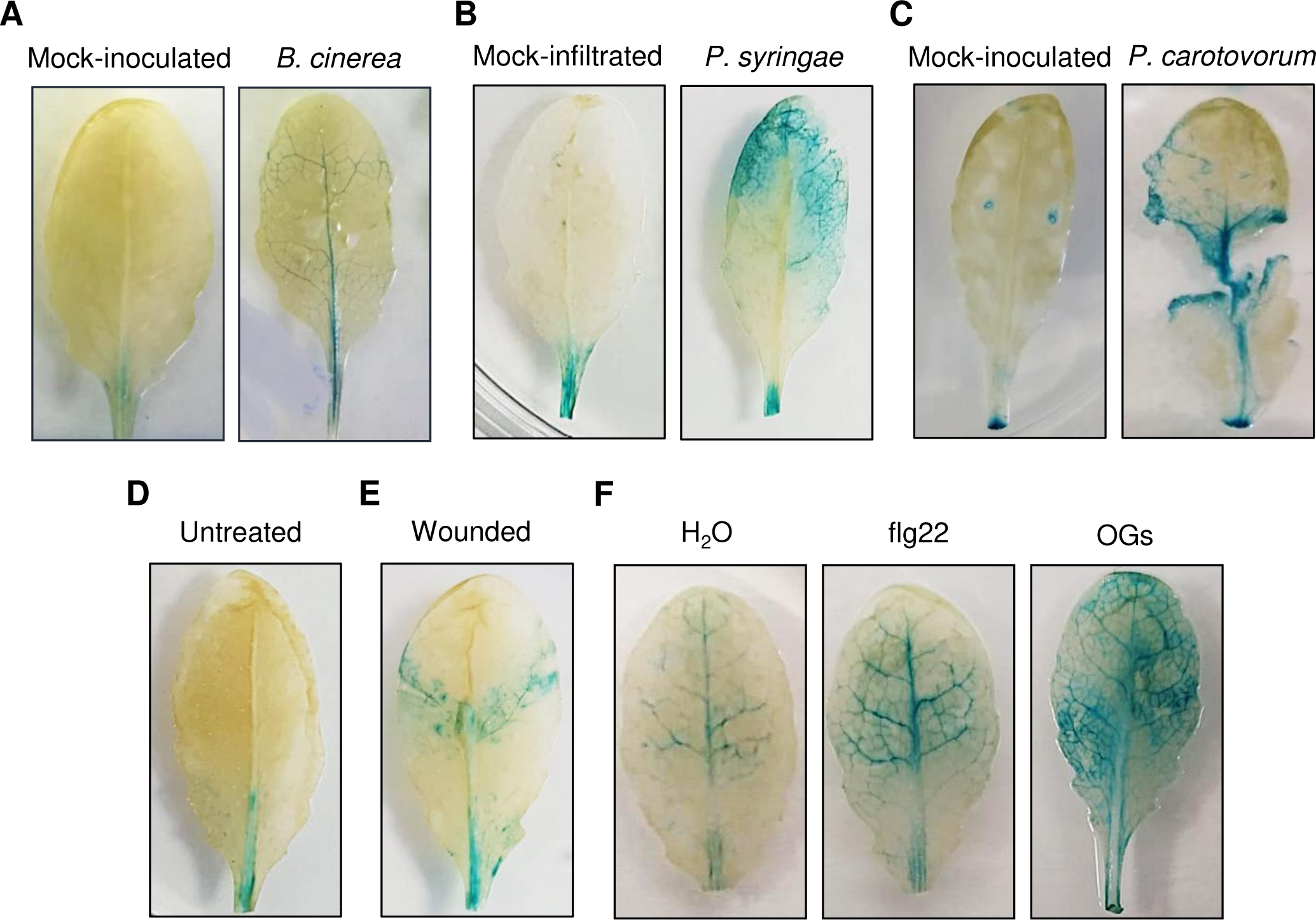
OGOX1::GUS transgenic leaves show increased GUS activity in response to pathogens, wounding and elicitor treatment. GUS activity was analyzed in OGOX1::GUS # 3.2 line. Results with a second independent line ((#2.3) are in Figure S1. **A.** Rosette leaves were drop-inoculated with *B. cinerea* conidia (5 μl, x10^5^spore/ml) or PDB (mock) as a control and GUS assay was performed 48 hours post- inoculation. **B.** Rosette leaves were infiltrated with *P. syringae pv. tomato (Pst)* DC3000 at OD=0.002 or with H2O (mock-infiltrated). GUS assay was performed 72 hpi. **C.** Rosette leaves were drop-inoculated at punctured sites with *P. carotovorum* cells (5 μl, OD600=0.025) or 50 mM potassium phosphate buffer pH 7.0 (mock-inoculated). GUS assay was performed 14 hpi. **D.** Untreated excised leaves analyzed at 72 h as a control. **E.** Rosette leaves were wounded by crushing with knurled tweezers; GUS assay was performed 1 hour after crushing. **F.** Rosette leaves were infiltrated with OGs (200 μg/ml), flg22 (100 nM) and water as control. GUS assay was performed 1 hour post-infiltration.

### The expression of OGOX affects the levels of OGs and oxidized OGs (oxOGs) *in planta*

The impact *in planta* of an altered oxidation of OGs was studied in available over-expressing lines of *OGOX1* (OGOX1-OE #1.9 and #11.8) (*18*) and loss-of-function mutants, obtained in this work. In both overexpressing lines compared to the wild type, transcripts of *OGOX1* were about 10 and 30 times higher, respectively (*18*). Besides *OGOX1*, also *OGOX2* shows expression in leaves [fig. S2, data are from eFBrowser (*23*)], which is not inducible during the immune response (*17*). A contribution of *OGOX2* in concert with *OGOX1* to the homeostasis of OGs in leaves cannot, therefore, be excluded and we decided to generate a *ogox1/2* double loss-of-function mutant. As the two genes are closely linked on chromosome 4 (*17*) and cannot be separated by genetic crossing, double loss-of-function mutants were obtained by CRISPR/Cas9 genome editing (fig. S3). Two *ogox1/2* double mutant lines (#1.5 and #4.6) were selected carrying frameshift mutations in both *OGOX1* and *OGOX2* in the homozygous state (Fig. 3).

**Fig. 3.**
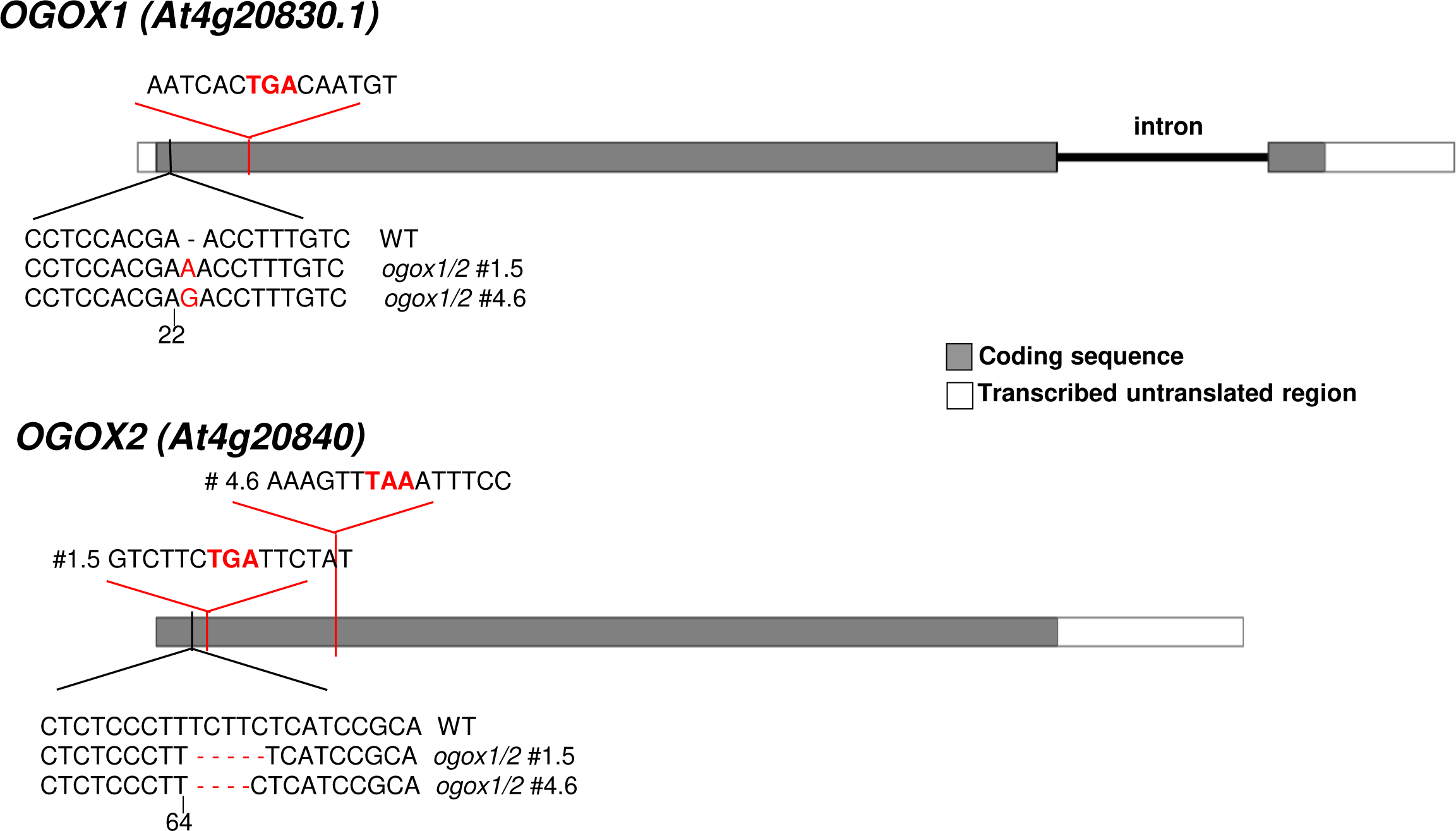
Schematic representation of the position of the frameshift mutations introduced by gene editing in both *OGOX1* and *OGOX2* in the double mutants *ogox1/2* lines #1.5 and #4.6. The single-nucleotide insertions after nucleotide 22 in *OGOX1* and the four- or five-nucleotide deletions after nucleotide 64 are all frameshift mutations that introduce stop codons (shown above in bold red).

How oxidation affects the fate of both exogenous and endogenous OGs in adult leaves of the overexpressing lines and the double *ogox1/2* mutants was assessed. Levels of OGs and their oxidized counterpart (oxOGs) were determined by High-Performance Anion-Exchange Chromatography (HPAEC-PAD) in the fraction indicated as Chelating Agent Soluble Solid (ChASS) from total CW preparations [Alcohol Insoluble Solids (AIS)] of leaves from WT, OGOX1-OE and *ogox1/2* double mutants infiltrated with OGs or water (*24*). Upon infiltration with OGs, higher levels of OGs and, conversely, lower levels of oxOGs were detected in the *ogox1/2* double mutants as compared with the WT. The opposite was found in the OGOX1-OE plants (Fig. 4A), indicating that upon infiltration, levels of OGOXs influence the relative balance between active OGs and oxOGs.

**Fig. 4.**
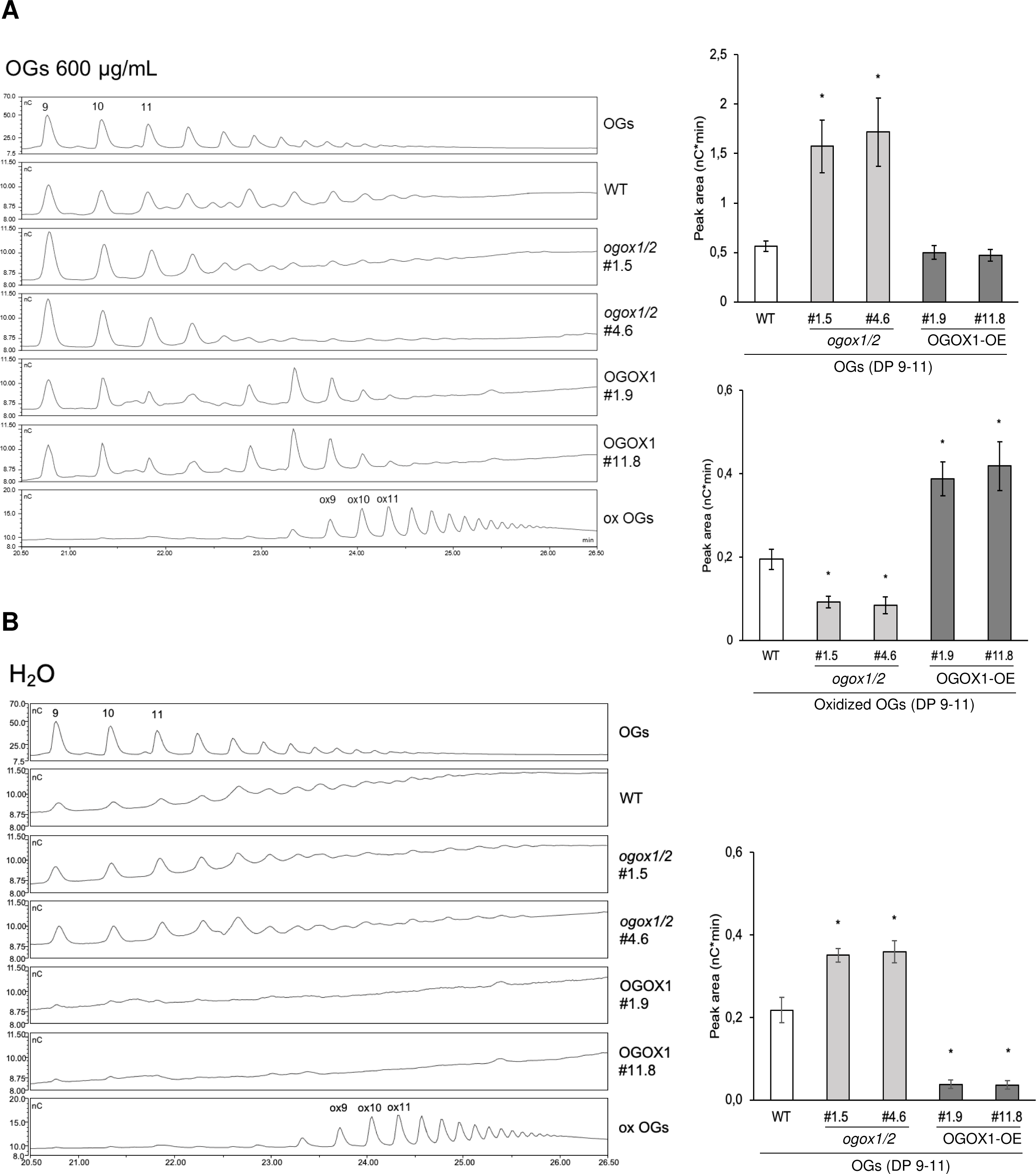
HPAEC-PAD analyses of cell wall chelating agent-extracted oligosaccharides from leaves of WT, *ogox1/2* and OGOX1-OE plants. Leaves were infiltrated with (**A**) OGs (600 μg/mL) or water (**B**) and the fraction containing OGs (Ch2SS) obtained from total cell wall preparations (AIS). Chromatographic profiles are shown on the left with OGs and oxidized OGs indicated by numbers corresponding to the degree of polymerization (DP). Graphs on the right show the sum of peak areas of OGs and oxidized OGs DP=9-11 in the chromatographic profiles. No oxidized OGs were detected in water-infiltrated leaves. An oxidized OG preparation of DP 9-16 was used as a standard.

Remarkably, in the water-infiltrated leaves, we detected i) higher levels of OGs in the *ogox1/2* mutants as compared with the wild-type, ii) lower levels of OGs in the overexpressing plants, iii) no oxOGs in both OGOX1-OE and *ogox1/2* plants (Fig. 4B). Overall, these results show that the level of OGs is determined *in planta* by levels of expressed OGOX.

### Expression of OGOXs affects defense-related gene expression and callose deposition induced by OGs

Whether signaling by OGs is affected by their enzymatic oxidation was further investigated by analyzing gene expression in leaves of plants altered in OGOX levels, upon infiltration with OGs or water as a control. The following defense-related genes as readouts of the OG signaling were examined: *FLG22-INDUCED RECEPTOR-LIKE KINASE 1* (*FRK1*), *CYP81F2*, *RESPIRATORY BURST OXIDASE HOMOLOGUE D* (*RBOHD*), analyzed at 1 h post-infiltration, and *PHYTOALEXIN DEFICIENT 3* (*PAD3*) analyzed at 3 hpi (*17, 25, 26*). Expression of all four marker genes was not significantly different in the OGOX1-OE leaves, compared to the wild-type, upon infiltration with water, whereas was lower upon infiltration with OGs (Fig. 5). This behavior is expected if the levels of active OGs decrease due to the enzyme action. In the *ogox1/2* mutants compared to the wild-type, expression of the four marker genes was differentially affected upon infiltration with OGs. While levels of *FRK1* and *PAD3* were higher in the OG-infiltrated leaves, in agreement with the higher level of active OGs expected in the mutants, expression of *RBOHD* and *CYP81F2* was lower. No significant difference in the expression of all marker genes was observed in water-infiltrated leaves compared to the wild-type. The differential expression behavior of the genes in the *ogox1/2* mutants suggests that factors other than the mere balance between active OGs and inactive oxOGs influence the complex gene expression network that regulates immunity.

**Fig. 5.**
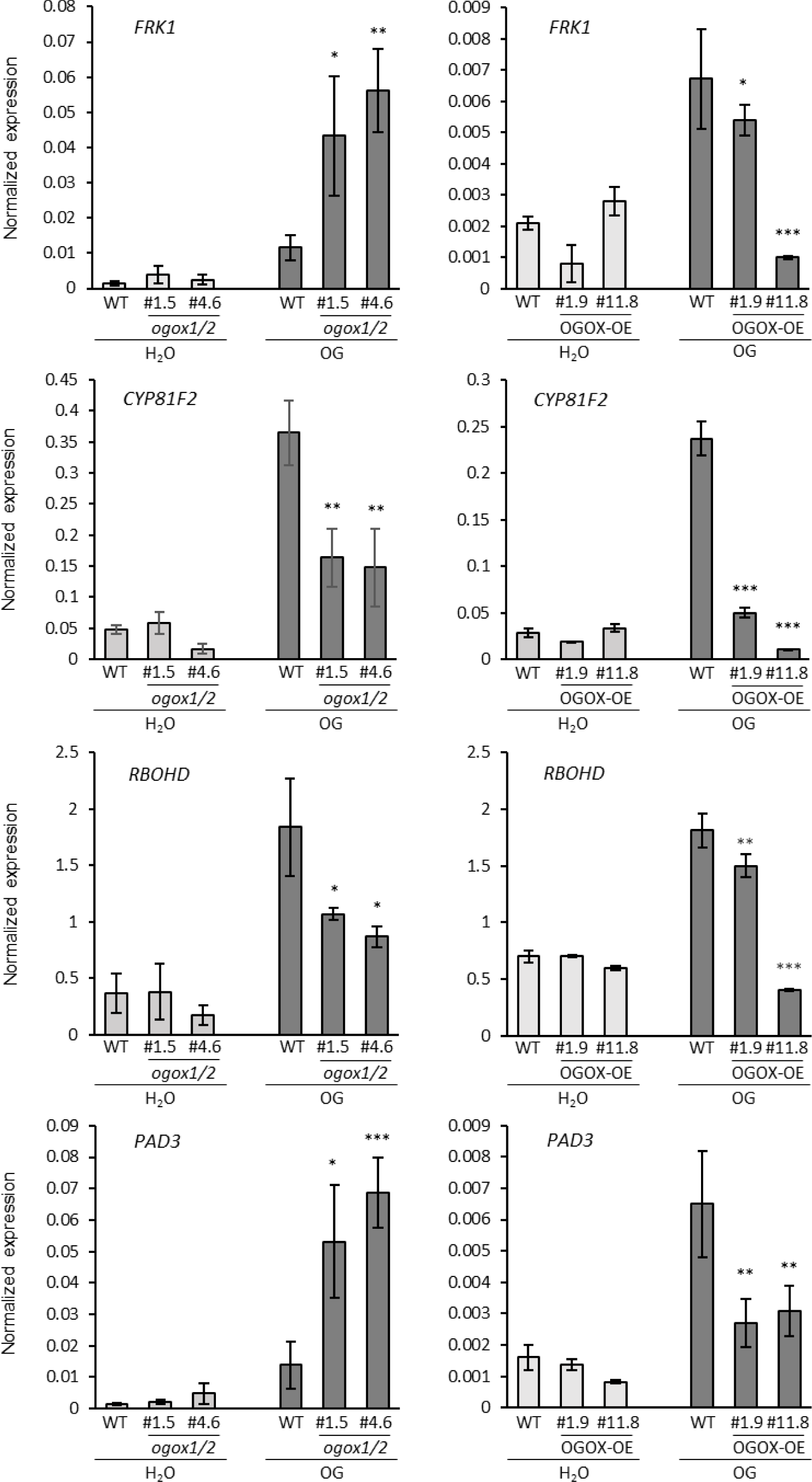
Plants altered in OGOX expression show altered expression of defense-related marker gene in response to OGs. Expression was analyzed by quantitative RT-PCR in rosette leaves from four-week-old plants of *Arabidopsis* Col-0 WT, OGOX1-OE overexpressing plants (lines #1.9 and #11.8), and CRISPR Cas-deleted *ogox1/ogox2* plants (lines #1.5 and #4.6) at 1 h (*FRK1, CYP81F2, RBOHD*) and 3 h (*PAD3*) post-infiltration with OGs (60 µg/ml) or water as control. *UBQ5* transcript levels were used for normalization. The mean of three biological replicates (± SD) is shown. Asterisks indicate statistically significant differences of mutants compared to WT treated plants according to Student’s t-test. (* p < 0.05; ** p < 0.01; *** p < 0.001).

Next, we investigated how alterations of OGOX expression affect a well-known late response to elicitors, namely callose deposition (*27*). Fluorescence microscopy images (Fig. 6A) of OG- infiltrated leaves, compared to the WT, showed a lower deposition of callose after 18 h in the *ogox1*/2 mutants and a higher callose deposition in OGOX1-OE lines. This result supports the view that not all the alterations observed in the OGOX manipulated plants can be interpreted in terms of levels of active OGs. Indeed, callose deposition has been shown to depend on the apoplastic H2O2 induced early after elicitation. For example, the lack of callose deposition observed in mutants lacking the apoplastic peroxidase PRX34 and the consequent apoplastic burst is rescued by adding H2O2 (*28*). The action of OGOX, on the one hand, decreases the levels of elicitor-active OGs but, on the other hand, increases the levels of H2O2 in the apoplast in the presence of the OGs as a substrate. It is likely, therefore, that the H2O2 generated by OGOX is responsible for the higher callose deposition in OGOX1-OE lines. Indeed, OGOX1-OE seedlings showed an increased H2O2 production compared to WT, whereas a significantly lower level of H2O2 was detected with the *ogox1/2* mutants. This may depend on the rapid action of OGOX present in the tissues on the added OGs (Fig. 6B).

**Fig. 6.**
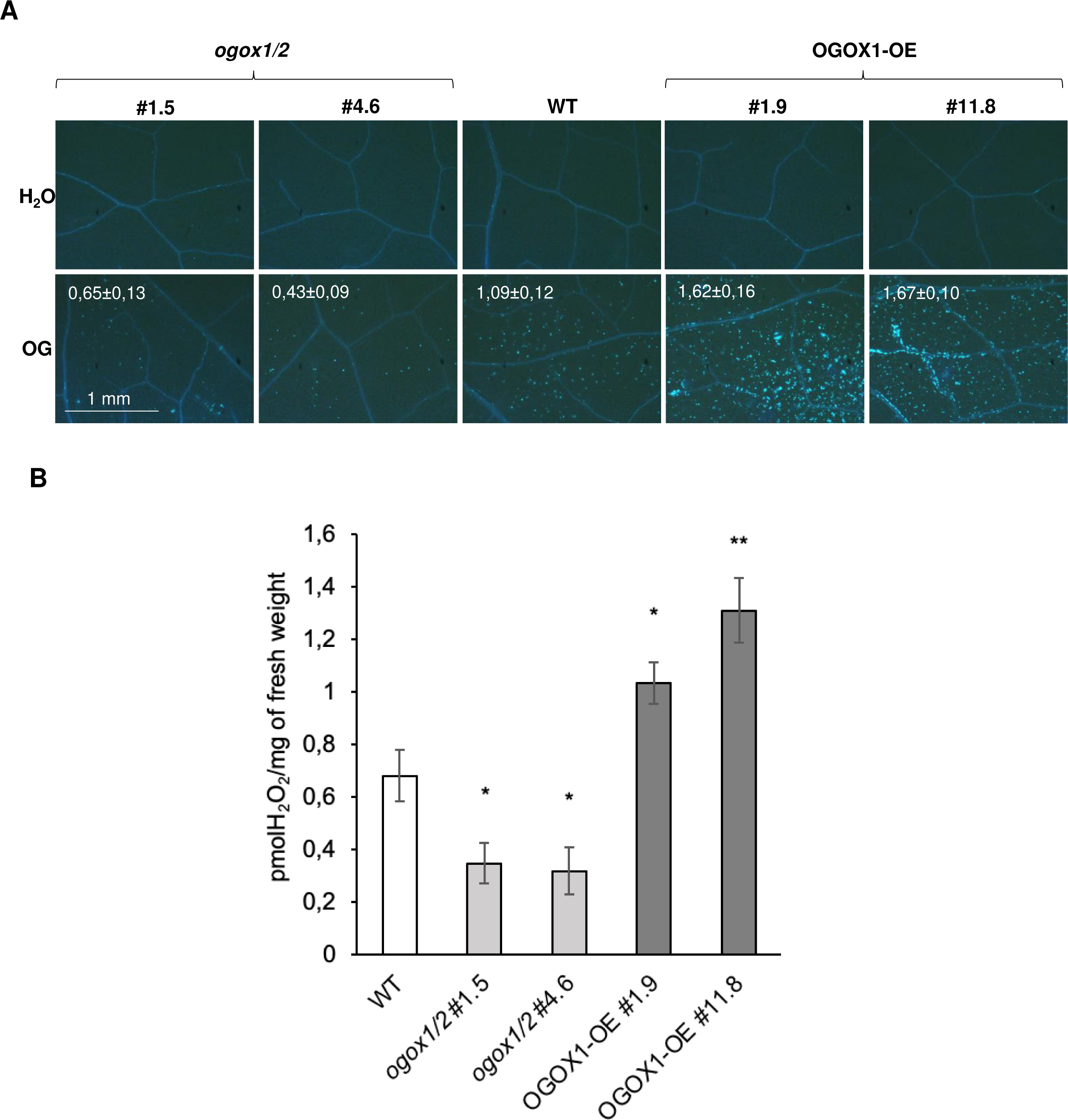
Responses of WT and OGOX-related mutants after treatment with OGs. **A.** Callose deposition in WT, *ogox1/2* and OGOX1-OE leaves after infiltration with OGs and H2O. Callose deposits were stained with aniline blue 24 h after treatment. Each picture is representative of at least 18 pictures acquired per genotype per treatment. Callose deposits were counted using the “analyze particles” function of ImageJ. Values indicate fluorescent area/background ± SE. Values for callose deposition of *ogox1/2* and OGOX1-OE lines were significantly different from WT, with a p value < 0.05 as determined by Student’s t-test.. **B.** OG-induced H2O2 accumulation in the culture medium of WT, *ogox1/2* and OGOX1-OE seedlings. Measurements were taken 30 minutes after treatment with 30 μg/mL OGs. Values represent the mean of at least 6 replicates ± Standard Error (SE). Asterisks indicate statistically significant differences of mutants compared to WT according to Student’s t-test. (n.s., not significant; * p < 0.05; ** p < 0.01).

Collectively, these data indicate that the alterations of OGOXs levels affect the complex interactions between the signaling activity of OGs, H2O2, and possibly other unidentified factors that may modulate the plant defense.

### Basal resistance to pathogens is bent on the dynamics of OGs levels, their oxidation and the microbial capacity of degrading oxOGs

The above results clearly show that OGOXs have an impact on the signaling activity of OGs, which consequently may have an impact on the plant response to pathogens. To investigate this point, the response of plants altered in *OGOX* expression was monitored upon infection with the bacteria *Pst* and *Pc* and the fungus *Bc.* Lesions, measured at 16 and 72 hours upon inoculation with either *Pst* or *Pc,* respectively, were smaller in the *ogox1/2* mutants and larger in the OGOX1-OE lines compared to the WT indicating that oxidation of OGs reduces the plant resistance to these bacterial pathogens (Fig. 7A and 7B). Instead, the *ogox1/2* mutants were more susceptible (Fig. 7C) and the OGOX1- OE plants were more resistant to *Bc* because this fungus has a reduced capability of utilizing the oxOGs as a carbon source (*18*). OGOXs therefore appear to play a role as resistance factors in the interaction with *Bc* by converting an important carbon source such as OGs into less digestible products and determining a critical retardation of the microbial growth in the plant tissue. On the other hand, the increased susceptibility of the *ogox1/2* mutants to *Bc* suggests that the signaling activity of OGs in this interaction, unlike in the case of *Pst* and *Pc,* is marginal with respect to their role as a carbon source.

**Fig. 7.**
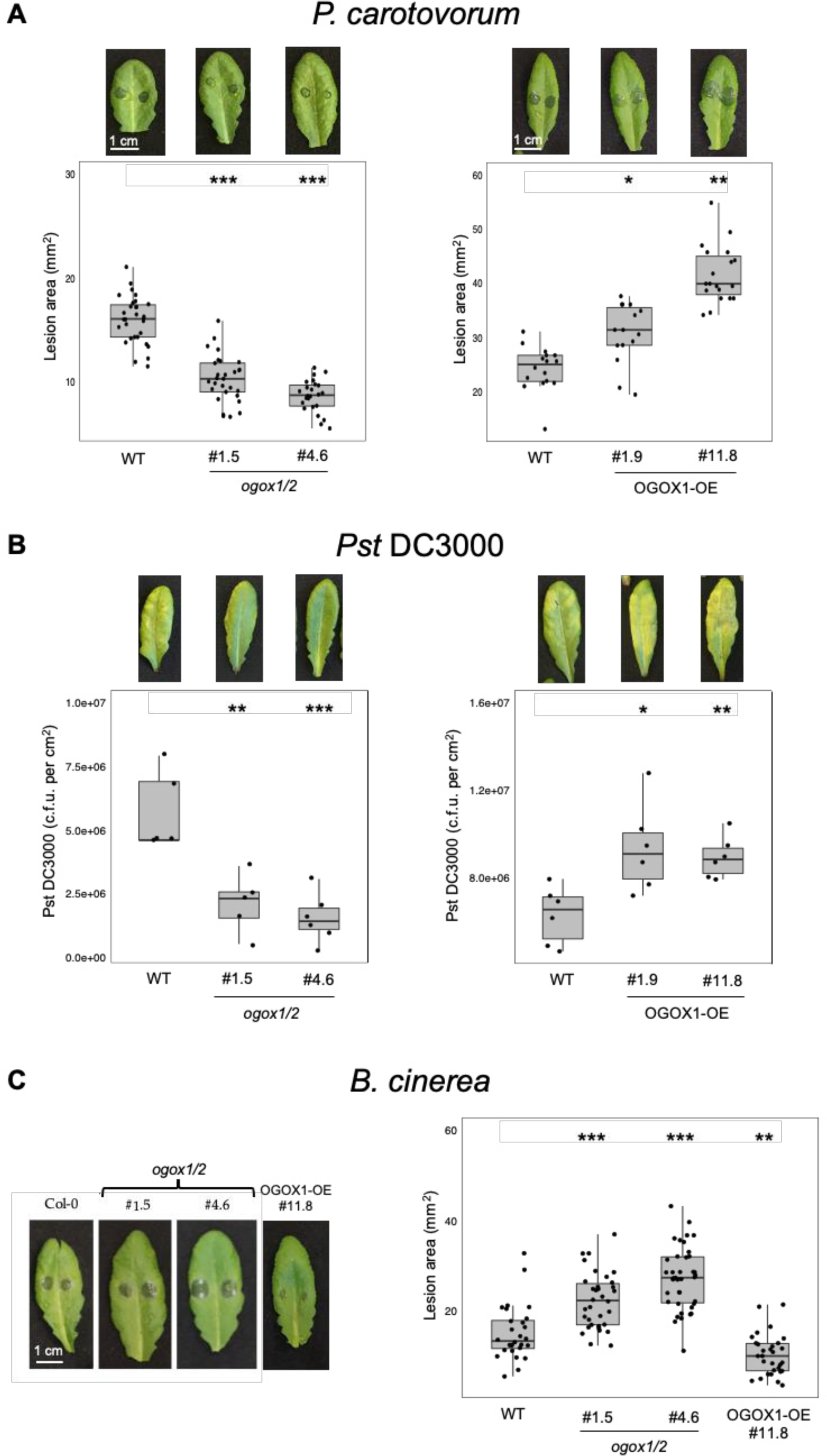
Plants altered in OGOX expression show altered response to pathogens. (A) *P. carotovorum,* (B) *Pst* DC3000, (C) *B. cinerea.* Lesion areas produced by *P. carotovorum* and *B.cinerea* were quantified at 16 and 48 hpi, respectively, using the ImageJ software. *Pst* DC3000 spread was quantified 72 hpi. Asterisks indicate statistically significant differences of mutants compared to WT according to Student’s t-test (* p < 0.05; ** p < 0.01; *** p < 0.001).

The utilization of oxOGs by *Pst* and *Pc* as a carbon source was examined in the current work by growing both bacteria, together with *Bc* as a control, in a minimal medium containing OG or oxOGs, respectively. Unlike *Bc*, which confirmed a reduced growth in the medium supplied with oxOGs, both *Pc* and *Pst* grew similarly in the media supplied either with oxOGs or OGs, indicating that both bacteria successfully metabolize both types of oligosaccharides as a carbon source (Fig. 8). We concluded that the increased resistance of *ogox1/2* mutants and increased susceptibility of OGOX1-OE plants to *Pc* and *Pst* depends on the relative amounts of elicitor-active OGs *vs.* inactive oxOGs, unlike in the case of *Bc*.

**Fig. 8.**
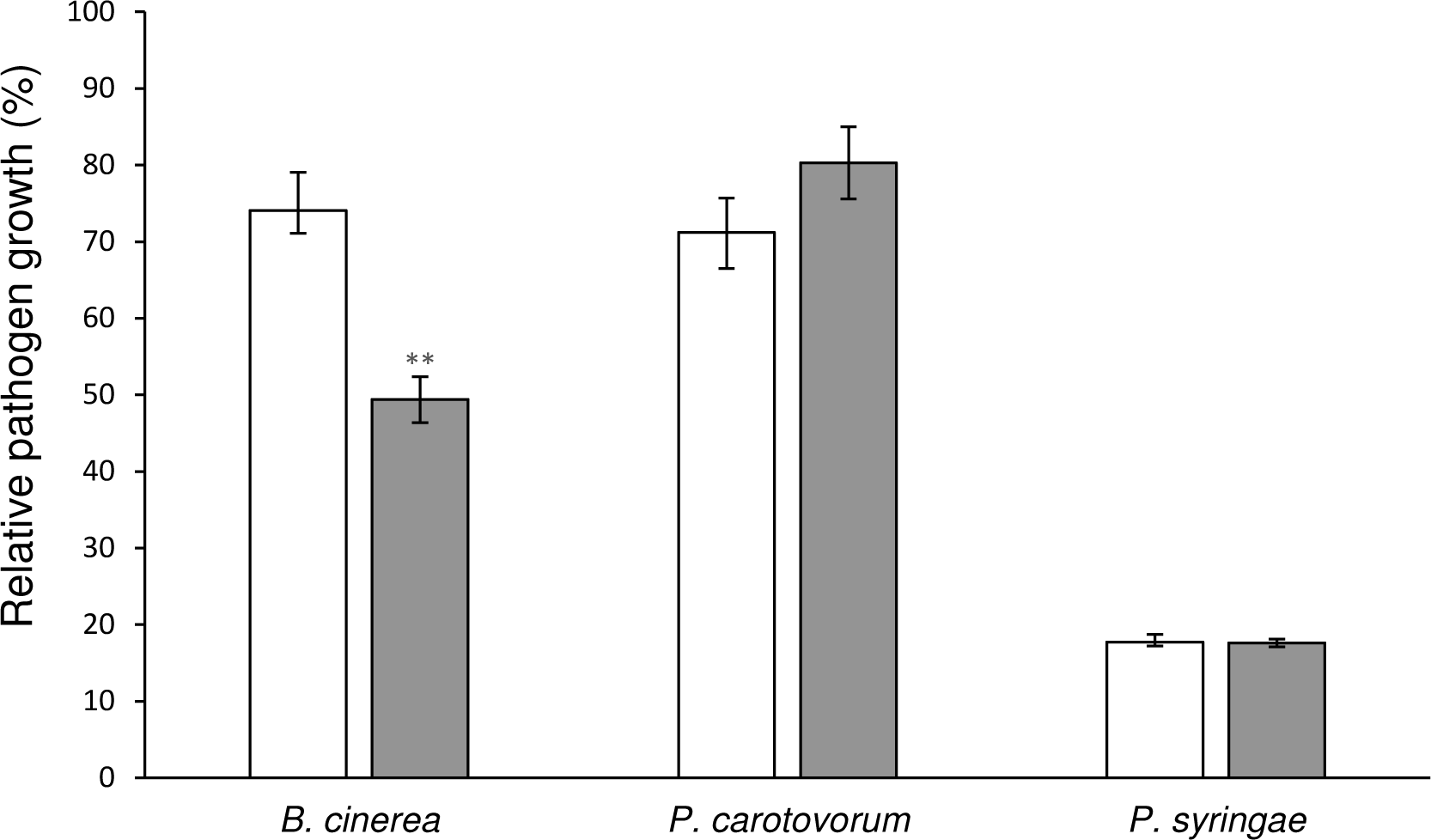
OG-to-growth conversion by *B. cinerea*, *P. carotovorum* and *P. syringae*. Relative growth of *B. cinerea*, *P. carotovorum* and *P. syringae* DC3000 in minimal medium supplemented with 0.15% (w/v) OG (white bars) and 0.15% (w/v) ox-OGs (grey bars) after 24 hour of incubation. Values are the ratio to the maximal growth observed in the same minimal medium supplemented with 0.15% (w/v) D-glucose. Values are mean ± SD (N=3). Asterisks indicate statistically significant differences according to Student’s t test (**, *p* < 0.005).

Since our results do not allow to distinguish the individual role of *OGOX1* and *OGOX2* in the response to pathogens, we also analyzed the contribution of *OGOX1* alone to the response to pathogens. A knock-out T-DNA insertional *ogox1* mutant was obtained that did not show detectable transcript levels and was therefore a null mutant (see characterization on fig. S4). Compared to the wild-type, the mutant showed increased susceptibility to *Bc,* similar to what observed in the *ogox1/2* mutants, whereas it showed no significant difference in its response to *Pst* and *Pc* (fig. S5A). These results indicate that only the simultaneous loss of *OGOX1* and *OGOX2* makes the plants more resistant to bacterial infection and suggest a redundant action of the two genes as susceptibility factors against bacterial pathogens, likely decreasing the levels of OGs in the infected tissues.

### Wound-induced accumulation of ROS increases in OGOX1 loss-of-function mutants

OGs have been proposed to act as local signals in the response to wounding (*7, 29–32*) and, indeed, in this work we show that *OGOX1* is transcriptionally up-regulated around a wound site (fig. 2E and fig. S1F). We therefore investigated whether OGOXs play a role in the response to a tissue damage by measuring the hydrogen peroxide generated at wound sites and in distal leaf veins as early as 1 hour after wounding (*33*). Hydrogen peroxide production at wound sites detected by DAB staining was increased in the *ogox1/2* mutants (fig. 9) as well as in the *ogox1* single mutants (fig. S5B) compared to wild-type plants.

**Fig. 9.**
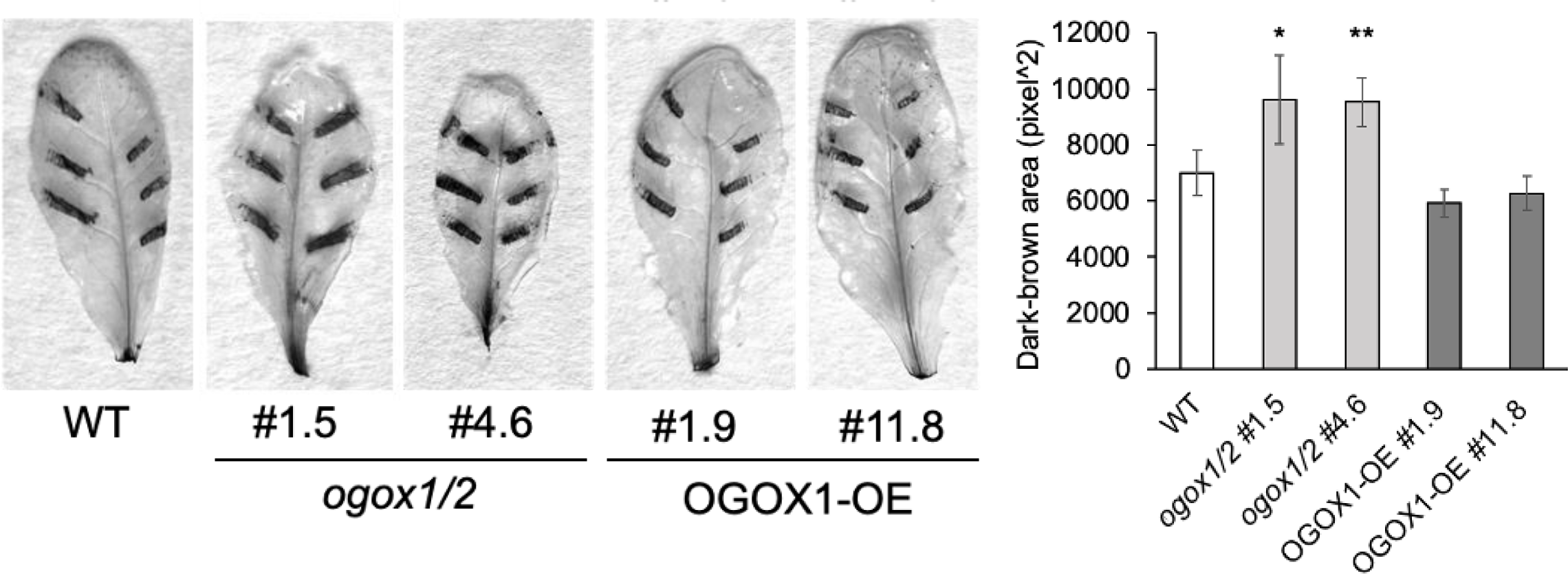
Wound-triggered hydrogen peroxide production is higher in the *ogox1/2* mutants. Leaves of four-week-old plants were wounded with knurled-tip tweezers, excised after 1 h and subjected to DAB staining for 4 h. *In situ* dark-brown precipitate generated by DAB oxidation in the presence of hydrogen peroxide was quantified using the “analyze particles” function of ImageJ. Dark-brown spots not generated by the tweezers’ wound, such as those at the excision site or in the central vein, were not considered. Values are the mean ± SD of at least three wounded leaves from three independent plants. Experiment was repeated three times with similar results. Asterisks indicate statistically significant differences of mutants compared to WT according to Student’s t-test (* p < 0.05; ** p < 0.01).

## DISCUSSION

We show in this work that OGOXs regulate *in planta* the levels of OGs acting as DAMPs and influence their immunity-related signaling and response to pathogens. Previous work demonstrated that overexpression of OGOX1 and in a similar way overexpression of CELLOX, which act on cellulose fragments and mixed linked beta-glucans, make plants more resistant to *Bc*, remarkably because this fungus is unable to metabolize oxidized oligosaccharides as a carbon source (*16–18*). Instead, we show here that the immune response is indeed influenced by the levels of OGOX- regulated elicitor-active OGs when pathogens metabolize equally well both non-oxidized and oxidized OGs as a carbon source for their growth. This is the case of *Pst* and *Pc*. Indeed, both bacteria have reduced ability to infect the *ogox1/2* double mutants, where the DAMP activity of OGs is higher, and increased ability to infect OGOX1-overexpressing plants, where the DAMP activity is lower. Against these pathogens, both *OGOX1* and *OGOX2* appear to play a redundant role, as suggested by the wild-type-like behavior of the single mutant o*gox1*. Unlike *OGOX1, OGOX2* is not up-regulated during the immune response and the encoded protein has been found to be subjected to post- translational modifications (PTM) in etiolated seedlings, after the cleavage of the 26-amino acid (aa) signal peptide for translocation into the ER. A new N-terminus lacking the first amino acid of the mature protein (an alanine) was found, likely generated by endopeptidase cleavage (*34*). Whether this or other kinds of PTM occur and modulate OGOX2 activity during immunity has never been investigated.

Taken together, our observations reinforce the notion that OGOXs, and likely other BBE-l proteins, play an important role in governing the potential DAMP activity of plants during the immune response. They also strengthen the notion that OGs are important players in plant immunity.

The most obvious physiological role of OGOXs is the homeostatic control of OGs to prevent an excessive accumulation that provokes plant growth penalties. The homeostasis of OGs ensures the proper defense-growth trade off that maximizes the recovery of a plant attacked by a microbe (*14*). The requisites of OGOXs for such a role are evident in the finding that these enzymes are apoplastic proteins up-regulated at the transcriptional level during pathogen attack, wounding and elicitor treatment, as shown in this paper. Alterations of their expression not only alter the fate of exogenous OGs in the expected direction but also influence the levels of OGs in untreated plants.

The orchestration of the OG levels by OGOX is evident in our gene expression analyses of the mutant/transgenic plants elicited with OGs. In the overexpressing plants, gene expression is affected in the expected direction towards an increased oxidation/inactivation of OGs. Namely all the genes (*FRK1*, *CYP81F2* and *RBOHD* examined at 1 h and *PAD3* at 3 h) showed a decreased expression due to the enzymatic oxidation of OGs impairing their signaling capability. Instead, in the *ogox1/2* mutants, the marker genes had a differential behavior, with *FRK1* and *PAD3* showing the expected increased up-regulation, while the expression of *CYP81F2* and *RBOHD* decreased, thus suggesting an impact of OGOX1 activity on immune pathways more complex that the simple inactivation of OGs as DAMPs. Noteworthy, both *RBOHD* and *CYP81F2* play an important role in elicitor-induced callose deposition (*28, 35*), but our observations show no correlation between the transcript levels of these two genes and callose deposition. Transcripts were lower in *ogox1/2* plants in response to OGs and even lower in the OGOX1-OE, while callose deposition was defective in the *ogox1/2* plants but increased in the OGOX1-OE plants. Moreover, although callose deposition and expression of *RBOHD* and *CYP81F2* have been shown to be controlled by a peroxidase-mediated apoplastic oxidative burst that enhances the activity of RBOHD establishing a feed-forward loop for H2O2 accumulation (*28, 36*), in our assays we observed no correlation between regulation of *RBOHD* and *CYP81F2* and entity of the extracellular oxidative burst induced by the addition of OGs. Worth of note, this burst was lower in the *ogox1/2* plants and higher in the OGOX1-OE plants, thereby showing no correlation with the non-oxidated active OG levels expected to be present in these plants. This apparent inconsistency may depend on the fact that H2O2 is produced during oxidation of the OGs by the OGOXs and increases the levels measured in our experimental system.

Notably, upon wounding, and therefore in the absence of massive levels of exogenously added OGs, the *ogox1/2* and the *ogox1* mutants showed a higher accumulation of ROS at the injured site in agreement with the expected higher levels of OGs in these plants. The action of OGOXs, which are up-regulated around a wound site as shown by our analyses, may prevent an excessive DAMP activity of OGs, likely to allow the initiation of the repair process, and contribute to the wound response (*7, 32*). Taken together, our results show that OGOXs are important players in the response to both pathogens and wounding.

The importance of OGs in plant signaling is emerging not only in plant-microbe interactions but also in events related to growth and development (*11, 14, 37–40*). Indeed, the ability of OGs to antagonize auxin points to these molecules as key factors at the crossroad between development and defense (*11, 41*). Moreover, evidence suggests that OGs are sensed through multiple and partially redundant perception/transduction complexes (*25*), besides that including WALL-ASSOCIATED KINASE 1 (WAK1) (*42*) and possibly some of its paralogs (*43*), to make plants resilient to the loss of a key signalling mechanism. The multiplicity of the OG sensing systems as well as the occurrence of compensatory mechanisms, never investigated yet, may explain contradictory results obtained with mutants silenced or deleted in the entire WAK family (*44–46*). In this context it is worth mentioning the recent finding that de-esterified homogalacturonan fragments, i.e. OGs, determine a “global” signaling by inducing clustering of cell surface regulators such as BRASSINOSTEROID INSENSITIVE 1 (BRI1), involved in development, and the PRR FLAGELLIN-SENSITIVE 2 (FLS2), inward bending of plasma membrane curvatures and a massive endocytosis of the clusters (*47*). This response is initiated by the binding of RAPID ALKALINIZATION FACTOR (RALF) peptide to de-esterified pectin fragments, i.e. OGs, which then undergo phase separation and recruit FERONIA and its coreceptor LLG1 [LORELEI-LIKE glycosylphosphatidylinositol-anchored protein (GPI-AP) 1] into pectin-RALF-FER-LLG1 condensates. Notably, RALF-induced endocytosis is abolished in the presence of OGOX1 (*47*), being dependent on long OGs, in agreement with the demonstration that OGOXs have a weak activity on short OGs (*14*).

Furthermore, a recent study has illuminated a novel biological mechanism wherein the physical interplay between a fungal PG and a plant PGIP facilitates the generation of long OGs with DAMP activity (*10*). This underscores that the generation of OGs is not a random occurrence but rather the outcome of a precise biological adaptation evolved by the plant to facilitate their release from the cell wall. In this paper, we reinforce the notion that specific mechanisms govern the levels of OGs *in planta*, by demonstrating that OGs are subject to a homeostatic mechanism mediated by OGOX, to regulate their activity.

Thus, the signaling activity of OGs, as previously discussed in many papers (*2, 48*), increasingly emerges as “global” and relevant not only in immunity but also in growth and developmental processes. How the release of OGs is finely and differentially tuned by the plant cell in many physiological events and how their activity is orchestrated by OGOXs becomes a frontier topic in plant biology.

## MATERIALS AND METHODS

### Plant material and growth

*Arabidopsis thaliana* wild type (WT) ecotype Columbia (Col-0) seeds were purchased from Lehle Seeds (Round Rock, TX; USA). The generation of OGOX1 overexpressing lines (OGOX1-OE #1.9 and #11.8) has been described previously (*18*). The *ogox1* (WISCDSLOX432E05) T-DNA insertional line (*49*) was obtained from the European Arabidopsis Stock Center (NASC). Plants were grown in a growth chamber at 22 °C, 70% humidity, under irradiance of 100 μE·m−2·s−1 with a photoperiod of 12-h light/12-h dark.

### Generation of constructs for stable transformation of Arabidopsis and selection of transgenic plants

*Agrobacterium tumefaciens* strain GV3101 was transformed with recombinant plasmids by electroporation and used for stable transformation of Arabidopsis accession Col-0, performed by the floral dip method (*50*).

To generate the OGOX1::GUS plants, the predicted promoter sequence of *OGOX1* was obtained from the Arabidopsis Gene Regulatory Information Server (AGRIS; arabidopsis.med.ohio- state.edu/). A fragment corresponding to 3002 nucleotides upstream of the predicted translation codon start of *At4g20830* was amplified from Arabidopsis Col-0 genomic DNA using the *BamHI*-*POGOX1* Fwd and *BamHI*-*POGOX1* Rev primers (table S1). The promoter fragments were cloned in the binary vector pBI121 (Stratagene) by a two steps reaction: (i) the CaMV 35S sequence was excided using the HindIII and XbaI restriction enzymes; (ii) the amplified fragment was cloned using the BamHI restriction site of pBI121 upstream of the *uidA* gene. From 6 independent transformed plants, four T3 homozygous lines (OGOX1::GUS, lines #2, #3, #4 and #5) containing a single insertion carrying the OGOX1::GUS cassette were characterized for the levels of the transgene transcript (fig. S1). Two lines (#2 and #3) were selected for further analyses.

For genome editing of *OGOX1* and *OGOX2*, plasmids were assembled using Golden Gate modular cloning method (*51*). Five gRNA sequences for *OGOX1* and *OGOX2* (see table S2) were chosen using “*chopchop.rc.fas.harvard.edu*” program. To generate the sgRNA expression cassettes, DNA fragments containing the ‘EF’ backbone with 67 bp U6-26 terminator were amplified using primers flaked with BsaI restriction sites associated with Golden Gate compatible overhangs (see table S1). The amplicons were assembled with the U6-26 promoter (pICSL90002) in Level 1 vectors (see table S2) according to the Golden Gate protocol. Briefly, 0.02 pmoles of purified PCR products were mixed with the same molar amount of the corresponding Level 1 and Level 0 (pICSL90002) vectors, 0.5 µl of BpiI enzyme (10 U/µl, ThermoFisher), 0.5 µl of T4 ligase (400 U/ µl, NEB), 1.5 µl of CutSmart Buffer (NEB), 10X Bovin Serum Albumin (1.5 µl) and water in a total reaction volume of 15 µl. The reaction was carried out as following: 20 seconds at 37 °C, 25 cycles of [3 minutes at 37 °C / 4 minutes at 16 °C], 5 minutes at 50 °C and 5 minutes at 80 °C. The resulting Level 1 vectors were then assembled in Level M vector (pAGM8055) following the same Golden Gate protocol. Cas9 expressing cassette with RPS5a promoter, the Cas9 coding sequence and the E9 terminator, together with the FAST-Red selectable marker (pBCJJ348) was generated by Castel et al. (*52*). Combination of the two-Level M vectors containing the sgRNAs and the Cas9 expression cassettes was assembled in Level P pICSL4723 binary vector according to the Golden Gate protocol.

For selection of *OGOX1* and *OGOX2* FAST-Red/Cas9/sgRNAs genome edited plants, twelve T1 transformed seeds (screened for the presence of the Fast-Red selection marker by fluorescence stereoscopy) were germinated and grown (fig. S3). Mutation in both *OGOX1* and *OGOX2* were found in two plants, #1 and #4, (fig. S3). T2 Cas9-free seeds without red fluorescence were selected, and single plants genotyped to confirm the presence of the mutations seen in T1: T2 plants *ogox1/2* #1.5 and #4.6 showed frameshift mutations in homozygosis in both *OGOX1* and *OGOX2* (Fig. 3). T3 and T4 seeds were obtained and genotyped, confirming the presence of the mutations in the homozygous state.

### Agroinfiltration and plasmolysis

Colocalization analyses were performed by *Agrobacterium tumefaciens-*mediated transient expression in tobacco leaves. PGIP2-RFP (*53*), and pm–rk (stock # CD3-1007, http://www.bio.utk.edu/cellbiol/markers/) were used as a cell wall marker and as a plasma membrane marker, respectively. The 35S::OGOX1-GFP [OGOX1.2 (*18*) and 35S::OGOX2-GFP constructs, with both OGOX1 and OGOX2 tagged with GFP at the C-terminus, were generated using the Gateway System vector (Thermo Fisher) . The 35S::CELLOX1-GFP construct was obtained using the Gateway system vectors, pDONR221 (Invitrogen), and pK7FWG2 (V141), using eGFP as a C-terminal tag (*54*). The 35S::CELLOX2-GFP construct has been previously described (*16*).

The infiltration buffer [50 mM MES, pH 5.6, 2 mM sodium phosphate buffer, pH 7.0, 100 μM acetosyringone, 0.5 % glucose] was used to resuspend the transformed *A. tumefaciens* for inoculation. OD at 600 nm was measured to have a final value of 0.5 for each sample. For co-transformation, *Agrobacterium* cultures grown separately and mixed prior the infiltration of *N. tabacum* leaves using a needless syringe. The infiltrated leaves were left for 48 h in the growth chamber before confocal microscope visualization, using a Zeiss LSM 710 microscope. For plasmolysis experiments, leaf disks at 2 days post-infiltration were treated with 1 M NaCl, incubated for 10 minutes and observed at the confocal microscope as previously described (*55*).

### Gus Analyses

Histochemical staining for GUS activity was performed by incubating leaves in the staining buffer (0.5 mg/ml X-Glca [5-Bromo-4-chloro-3-indolyl β-D-glucuronic acid cyclohexylammonium salt; Duchefa Biochemie]; 2 mM K3[Fe(CN)6]; 2 mM K4[Fe(CN)6]; 0.2% Triton X-100; 50 mM buffer sodium phosphate, pH 7.2; 2% DMSO) overnight at 37 °C, with shaking. Subsequently, the samples were cleared and dehydrated with 100% boiling ethanol and rehydrated in 50% ethanol before photography.

For the response to elicitors, adult leaves were infiltrated with water, flg22 (100 nM) or OGs solution (200 μg/mL) using a needleless syringe and, after 1 h, were detached from plants and assay was performed. For the response to wounding, adult leaves were wounded by applying a single pressure on each side of the lamina flanking the middle vein with a laboratory forceps. After 1 h, leaves were detached from plants and assay was performed. Infections were performed as described below.

### Infection Assays

*Botrytis cinerea* (*Bc*) conidia were suspended in potato dextrose broth (PDB, 24 g/L; Difco, Detroit, USA) at a final concentration of 5 × 10^5^ conidia/mL and incubated for 2-3 h at room temperature before inoculation of Arabidopsis leaves previously described (*56*). Adult leaves were inoculated with 5-μl drops of a conidiospore suspension (5 x 10^5^/mL) in PDB or PDB only (mock), regularly spaced on each side of the middle vein and, after 48 h, leaves were detached for analysis (GUS staining or symptom analysis).

*Pectobacterium carotovorum* subsp. *carotovorum* strain DSMZ 30169 (*Pc*) was grown as previously described (*32*). For inoculation, adult leaves were punctured with the sterile needle of a 1-mL syringe on the epidermis of the adaxial surface of each leaf, at the sides of the middle vein, and a 5-μl droplet of the bacterial suspension (OD600=0.05) or mock (50 mM potassium-phosphate buffer, pH 7.0) was placed on each punctured site. Leaves were excised after 16 h for GUS staining. Symptoms caused by *Bc* and *Pc* were assessed by measuring the area of macerated tissue (lesion area), at 48 and 16 hours post-inoculation respectively, using ImageJ software. (https://imagej.nih.gov/ij/).

*Pseudomonas syringae* pv *tomato* (*Pst*) DC3000 was cultured in LB liquid medium containing 25 μg/mL rifampicin and inoculations were performed by infiltrating adult leaves with water or the bacterial suspension (5 x 10^4^ cfu/mL) using a needleless syringe as previously described (*57*). GUS staining was performed after 3 days. Bacterial growth was quantified at 0- and 3-days post infiltration.

### Preparation of OGs and oxidized OGs

The oligogalacturonide (OG) standard were prepared in vitro through the partial digestion of purified polygalacturonic acid (PGA) using endo-polygalacturonase II from *Aspergillus niger*, as outlined (*58*). The oxOGs were obtained by incubating the standard OGs with OGOX1 expressed in *Pichia pastoris*, as previously described (*18*).

### Analysis of OGs and oxidized OGs in Arabidopsis plants by HPAEC-PAD (High-Performance Anion-Exchange Chromatography with a Pulsed Amperometric Detector)

Two leaves (about 50 mg of fresh weight) of 4-week-old plants were infiltrated each by using a syringe without the needle, with sterile ultrapure water or with a solution containing 600 ng/μL OGs. Leaves were immediately excised, frozen in liquid nitrogen and homogenized for 2 min at 30 Hz in a mixer mill MM301 (RETSCH), using inox beads (6 mm diameter). To isolate cell wall polysaccharides that are Alcohol Insoluble Solids (AIS), ground tissues were re-suspended in 1 mL of 70% v/v ethanol pre-warmed to 70°C. The pellet was washed twice with a chloroform:methanol (1:1, vol/vol) mixture, vortexed, and centrifuged at 14000 g for 10 minutes. Following this, it was washed twice with acetone and centrifuged at 14000 g for 10 minutes. Finally, the pellet was dried at room temperature under a chemical hood overnight. For extraction of the Chelating Agent Soluble Solids (ChASS) fraction containing OGs, AIS fractions were re-suspended in 200 μl chelating solution 2 (ChA2: 50 mM CDTA, 50 mM ammonium oxalate, 50mM ammonium acetate, pH5.5) and incubated overnight at 4 °C as detailed (*24*). Polysaccharides obtained from the ChASS extraction were dissolved in ultrapure and were subsequently analyzed with HPAEC-PAD by using equipment and procedures as previously described (*18*).

### Gene expression analysis: RNA extraction, RT reaction and Real Time RT-PCR

Leaves of 4-week-old Arabidopsis plants were infiltrated in a single site with a solution containing OGs (60 μg/mL). Total RNA was extracted using RNA isolation NucleoZol (Macherey-Nagel) according to the manufacturer’s instructions and treated with RQ1 RNase-free DNase (Promega). cDNA was synthesized with ImProm-II™ Reverse Transcription System (Promega). qRT-PCR was performed with a CFX96 Real-Time PCR System (BioRad) using iTaq Universal SYBR Green Supermix (BioRad) as recommended by the manufacturer. The amplification protocol consisted of 30 s of initial denaturation at 95°C, followed by 45 cycles of 95°C for 15 s, 58°C for 15 s and 72°C for 15 s. Primers used for gene expression analysis are listed in table S1. The expression levels of each gene, relative to UBQ5, were determined using a modification of the Pfaffl method (*59*) as previously described (*60*).

### Callose deposition assays

Callose deposition was detected as previously described with some modifications (*61*). Leaves of 4- week-old plants were infiltrated with a solution containing OGs at a final concentration of 60 μg/mL; detection was performed 24 h post-infiltration. After staining, leaves were mounted in 50% (v/v) glycerol and examined by a UV epifluorescence microscope (Nikon, Eclipse E200) equipped with a cooled charge-coupled device camera (DS-Fi1C). Images were acquired with the Nis Elements AR software (Nikon). Callose deposits were counted using the “analyze particles” function of ImageJ.

### Xylenol Orange Assay

Extracellular H2O2 by OG-treated 13-days-old seedlings was measured by a colorimetric method (*62*) based on the peroxide-mediated oxidation of Fe^2+,^ followed by the reaction of Fe3^+^ with the xylenol orange dye (o-cresolsulfonephthalein 3′,3″-bis[methylimino] diacetic acid, sodium salt; Sigma). Ten seeds were placed in each well of 12-well plates containing 2 mL of liquid MS/2 medium. At least 9 biological replicates for genotype for treatment were set up. One day before treatment medium was refreshed and levelled off. The treatment has been performed adding an OG solution to the medium to reach a final OG concentration of 30 μg/mL. H2O were used as control. Thirty minutes post treatment 100 μL of medium was collected and 100 μL of xylenol orange final reagent solution was added. Standard curves of H2O2 were obtained for each independent experiment. Absorbance (A560) of the Fe^3+^-xylenol orange complex was detected by microplate reader (BioRad). Data were normalized and expressed as picomolar H2O2/mg fresh weight of seedlings.

### Pathogen growth assays on OGs and oxidized OGs

Growth assays of *B. cinerea* on OGs and oxidized OGs were done as previously described (*18*). *P. carotovorum* subsp. *carotovorum* and *P. syringae* DC3000 were grown in a minimal medium (13 mM potassium phosphate pH 7, 17 mM NaCl, 1.7 mM sodium citrate, 30 mM (NH4)2SO4 and 2.8 mM MgSO4) supplemented with the appropriate carbon source. Bacterial growth was carried out on a rotary shaker (160 rpm) at 28°C using 1,.4*10^7^ cell/mL as starting inoculum. Growth was determined spectrophotometrically by measuring the optical density at 600 nm (OD600) after 24 h of incubation. Three replicates for each growth were performed. For the growth of all three pathogens in the presence of OGs or oxOGs, the minimal medium was supplied with OGs or oxOGs with a degree of polymerization ranging from 4 to 10 at a final concentration of 0.15 % (w/v) before filter-sterilization. Pathogen growth (%) was calculated as the percentage ratio between the growth measured in presence of OGs or oxOGs and that in minimal medium supplemented with 0.15% (w/v) D-glucose.

### DAB staining upon wounding

H2O2 visualization in response to wounding have been performed basically as already described (*28*). Briefly, leaves of 4-week-old Arabidopsis plant were wounded by applying a single pressure on each side of the lamina flanking the middle vein with a laboratory forceps and, after 1 h, were detached from plants and dipped for 4 h in a solution containing 1 mg/mL 3,3- diaminobenzidine (DAB) pH

5.0 at room temperature, with shaking. Subsequently, the samples were cleared and dehydrated with 100% boiling ethanol and rehydrated in 50% ethanol before taking pictures.

## Acknowledgments

We thank Irene D. Romano for technical support during the experiments. We acknowledge the invaluable contributions of Professor Gabriella Piro, who sadly passed away during the preparation of this manuscript, in protein localization analysis using confocal microscopy.

## Author contributions

Conceptualization: G.D.L., D.P., FC. Methodology: A.S., A.D., D.P. and G.D.L. Investigation: A.S., A.D., F.L., M.G., G.G., D.P., M.B., G.P., S.C., M.D.C., M.I. and B.C. Formal analysis: A.S., A.D Visualization A.S., A.D., D.P and G.D.L. Supervision: FC, D.P. and G.D.L., Resources: G.D.L. Writing—original draft: A.S., A.D. and G.D.L. Writing—review and editing: A.S., A.D., F.C., D.P and G.D.L. Funding acquisition: G.D.L. and D.P.

## Data and materials availability

All data needed to evaluate the conclusions in the paper are present in the paper and/or the Supplementary Materials. Request for plant materials should be submitted to GDL

## Supplementary Materials

Fig. S1. Characterization of the POGOX1::GUS transgenic plants.

Fig. S2. OGOX2 (At4g20840) is expressed in leaves.

Fig. S3. Sequential steps followed for the selection of CRISPR-Cas9 edited single and double ogox1 and ogox2 mutants.

Fig. S4. Characterization of the T-DNA insertional mutants ogox1.

Fig. S5. Analysis of pathogens resistance and wound-triggered H2O2 production in the ogox1 null mutant and in the OGOX1-overexpressing lines.

Table S1. Primers used in this work.

Table S2. sgRNAs targeting sequences and Level 1 plasmids

